# Dandelion pappus morphing is actuated by radially patterned material swelling

**DOI:** 10.1101/2021.08.23.457337

**Authors:** Madeleine Seale, Annamaria Kiss, Simone Bovio, Ignazio Maria Viola, Enrico Mastropaolo, Arezki Boudaoud, Naomi Nakayama

## Abstract

Plants can generate motion by absorbing and releasing water. Such motion often facilitates reproductive success through the dispersal of seeds and fruits or their self-burial into the soil. Asteraceae plants, such as the dandelion, often have a hairy pappus attached to their fruits to allow them to fly and in many cases, these can open and close depending on moisture levels to modify dispersal. Here we demonstrate the relationship between structure, composition and function of the underlying hygroscopic actuator. By investigating the structure and properties of the actuator cell walls we have identified the mechanism by which the dandelion pappus closes and developed a structural computational model that can capture observed pappus closing. This model was used to explore the contribution of differential material domains in the actuator function and the critical design features. We find that the actuator relies on the radial arrangement of vascular bundles and surrounding tissues including cortex and the floral podium around a central cavity. This allows heterogeneous swelling in a radially symmetric manner to co-ordinate precise movements of the pappus hairs attached at the upper flank. This actuator is a derivative of bilayer structures, but is radial and can synchronise the movement of a planar or lateral attachment. The simple, material-based mechanism presents a promising biomimetic potential in robotics and functional materials.

## Introduction

Movement of body parts are typically mediated by specialised hinge structures – actuators – in biological and engineered systems. Biological actuators consist of continuous structures, and differential expansion within the actuator drives the reversible movement and morphing. A thematic example is bilayer structure, in which two sides of a planar or cylindrical body expands or shrink more to cause bending twisting. Inspired by the plethora of examples from nature, diverse designs have been developed for bilayer soft robotic actuators. Hygroscopic plant movements have been particularly relevant for biomimetic engineering and design as some of them do not rely on inherently biologically active processes^1^. Instead, they can highlight structural features that have been tuned by evolution to optimise mechanical efficiency or use of materials.

Plant movements and morphing are generally driven by changes in hydration^2^. This can be actively regulated by altering solute concentrations to manipulate osmotic gradients or by increasing water uptake and the prevalence of aquaporins^2,3^. Active water movement occurs in the opening and closing of stomata and the leaf curling of *Mimosa* plants^4,5^. Similarly, turgor pressure can allow rapid movements by exploiting mechanical instabilities of precisely formed tissues such as in the Venus fly trap and in the explosive dispersal of *Cardamine hirsuta*^6,7^. Alternatively, plant cell walls can passively absorb and release water to cause morphology changes^8,9^. These hygroscopic movements have been demonstrated, for example, in pine cones, wheat awns and ice plant seed capsules^10–12^.

Directed hygroscopic movements often arise from the differential expansion of cells within a tissue or parts of cell walls with different material properties. These materials respond to water in different ways to allow, for instance, anisotropic swelling typically resulting in bending or coiling motions^8^. For example, adjacent tissue types with alternating cellulose microfibril orientations generate a bilayer structure to cause bending or twisting motions^13–15^. This can be combined with differential deposition of phenolics. In the curling stems of the resurrection plant, *Selaginella lepidophylla*, different amounts of lignin are deposited on each side of the stem with increased hydrophobicity and elastic modulus observed for tissues where lignin is present. The non-lignified side can therefore absorb more water and deform more easily allowing the plant to unfurl its stems when wet and initiate photosynthesis^16,17^. Similarly, *Erodium gruinum* awns exhibit differential deposition of phenolic compounds in distinct tissue parts affecting the rate of curling along the length of the awn^18^.

In addition to material composition, the geometry of cells and tissues can contribute to controlling hygroscopic movements. *S. lepidophylla*, cells that swell less tend to have thicker cell walls^16,17^. In the seed capsules of the ice plant, *Delosperma nakurense*, cells with a honeycomb structure expand anisotropically due to their elongated geometry and the arrangement of cell wall layers within them^12^. These examples illustrate that both structural and compositional features are combined to facilitate appropriate and efficient hygroscopic motions, but all rely on heterogeneity of adjacent materials.

The haired fruit of the common dandelion undergoes morphing to open or close its flight-enabling pappus^19–21^. When the hairs are drawn together and the pappus is closed, the fluid dynamics around the pappus are dramatically altered and the dispersal capacity is modified^22^. This allows the plant to tune dispersal by optimising timing and distances in response to environmental conditions. The dandelion pappus changes shape via a hygroscopic actuator at the apical plate of the achene (fruit) that swells on contact with water^19–21^.

In addition to hygroscopic absorption of water by cell walls in the apical plate, an alternative pappus morphing mechanism occurs in dandelion pappi relying on the cohesive properties of water droplets. Fine hairs that easily bend are particularly sensitive to the cohesion forces generated by water when it forms a contact point with the solid hairs^23^. Bending of dandelion pappus hairs has been observed before in response to water droplets and may be useful inspiration for engineering precision liquid handling devices^24,25^.

While the hygroscopic actuator function of the apical plate has been observed before, its mode of action remains unclear. This actuator is composed of distinct domains originating from the floral podium, vascular bundles and surrounding cortex tissue. We have found that it generates a sophisticated and precisely patterned radial geometry of at least four different tissue types, differential swelling of which enables the reversible angular movement of the pappus hairs. This is more complex than previously described hygroscopic plant actuators which typically rely on one or two tissue layers in planar or cylindrical structures to generate bending or coiling. Radial swelling of the actuator allows the hairs to be pushed both outwards and upwards in contrast to a previous hypothesis suggesting that the hairs are pushed upwards via a lever-like mechanism^20^. Unlike other hygroscopic plant movements, the dandelion makes use of radially symmetric swelling to generate torque, which may help to synchronise movement of the roughly 100 hairs that are organised in a disk like geometry^26^.

## Results

### An actuator at the base of the pappus drives morphing

To investigate the mechanism and dynamics of reversible pappus closure, we imaged dandelion pappi in a bespoke hydration chamber^22^. The extent of pappus closure depends on the amount of water added to the chamber and the pappus reaches a steady state over a period of 30 – 60 minutes depending on the dynamics of water addition^22^. In our experiments the pappus angle typically changes by 40 – 100°.

We examined the apical plate structure at the base of the pappus where the hairs attach that is required for pappus closure and reopening (Fig 1h). We applied a resin to different parts of the apical plate expecting it to block the ability of the structure to move or swell (Fig 1a-b,d-e,g, Fig S1). It is possible that water entry into the tissue was also affected. In case (B), the resin was applied to the upper side of the apical plate, in (C) to the lower side and a control set (A) were left unchanged (Fig S1). Blocking the upper side of the apical plate (B) partially prevented pappus closure but blocking the lower side (C) almost completely prevented closure (Fig 1a-b,d-e,g). This indicated that pappus closure does not originate from bending of the hairs themselves, but is largely dependent on the central actuator and particularly on the lower side of it.

**Figure 1.**
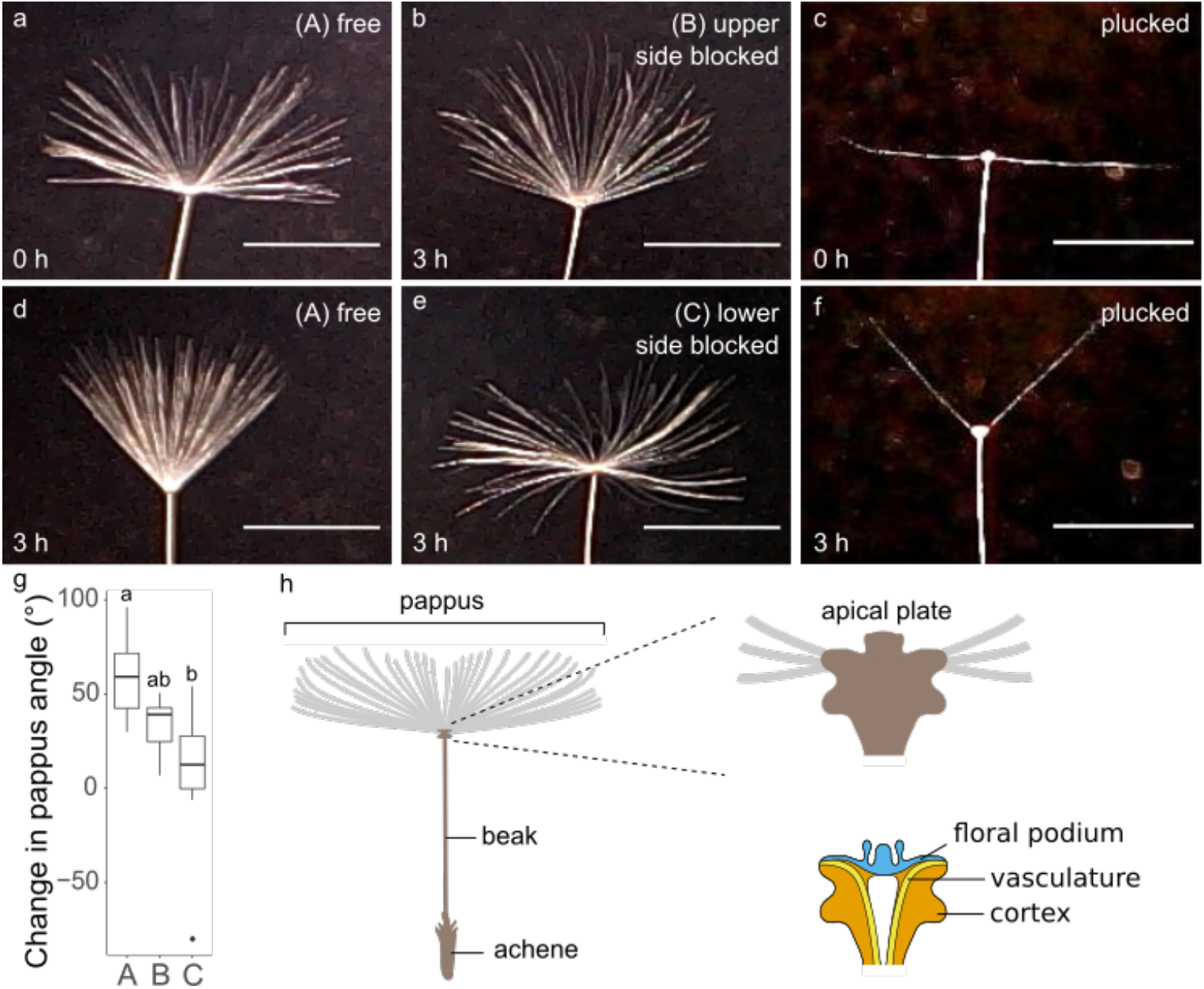
The apical plate is required for pappus morphing. **a-b,d-e** apical plate blocking experiments showing **a** a sample before hydration, and **b,d,e** 3 hours after wetting. **a** and **d** have no resin applied, **b** has resin applied to the upper side of the apical plate, **e** has resin applied to the lower side of the apical plate. Scale bars are 5 mm. **c,f** effect of removing hairs on pappus morphing. **c** a sample before hydration with all but two opposite hairs removed, **f** the same sample as in **c** 3 hours after hydration. **g** the change in pappus angle (between outermost hairs) between dry and wet conditions for apical plate blocking experiment, n = 10 per treatment. **h** illustrates the morphology of the diaspore, location of the apical plate, and a cross-sectional view of the apical plate indicating some internal structures.

**Figure S1.**
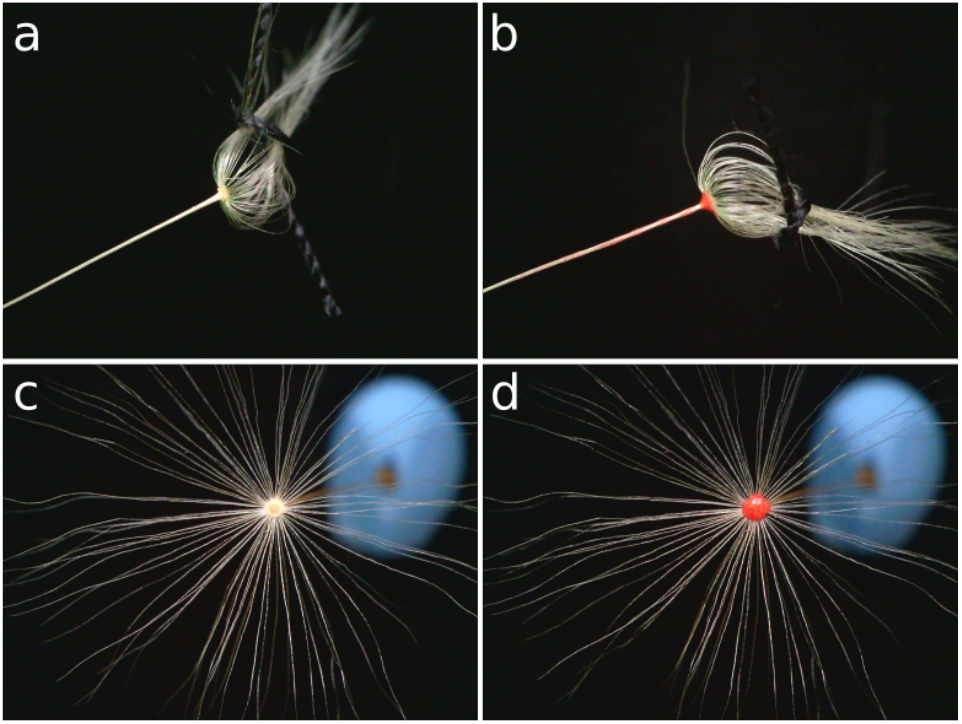
Images of the pappus illustrating methods of apical plate blocking. **a** tied pappus hairs before applying resin to lower side, **b** pappus after applying resin to lower side, **c** pappus before applying resin to upper side, **d** pappus after applying resin to upper side.

An alternative mechanism for pappus closure is via adhesion of the hairs to one another due to surface tension and capillarity. This has been previously demonstrated for dandelion pappi when they hold large droplets of water^24^. To ascertain the role of hair adhesion in the small droplet-derived closing observed here, we removed most of the hairs from dandelion pappi to massively increase the spacing between them (Fig 1c,f). This would prevent adhesion between neighbouring hairs as demonstrated by Bico *et al.*^23^. We found that dandelion pappi with just two hairs remaining were still able to close in response to moisture addition (Fig 1c,f, Fig S2). The dynamics and magnitude were in fact slightly enhanced compared to intact samples (Fig S2). This may be because clusters of hairs normally slightly obstruct one another during motion and removing hairs reduces this effect. These data indicate that surface tension is not involved in this type of pappus closing.

**Figure S2.**
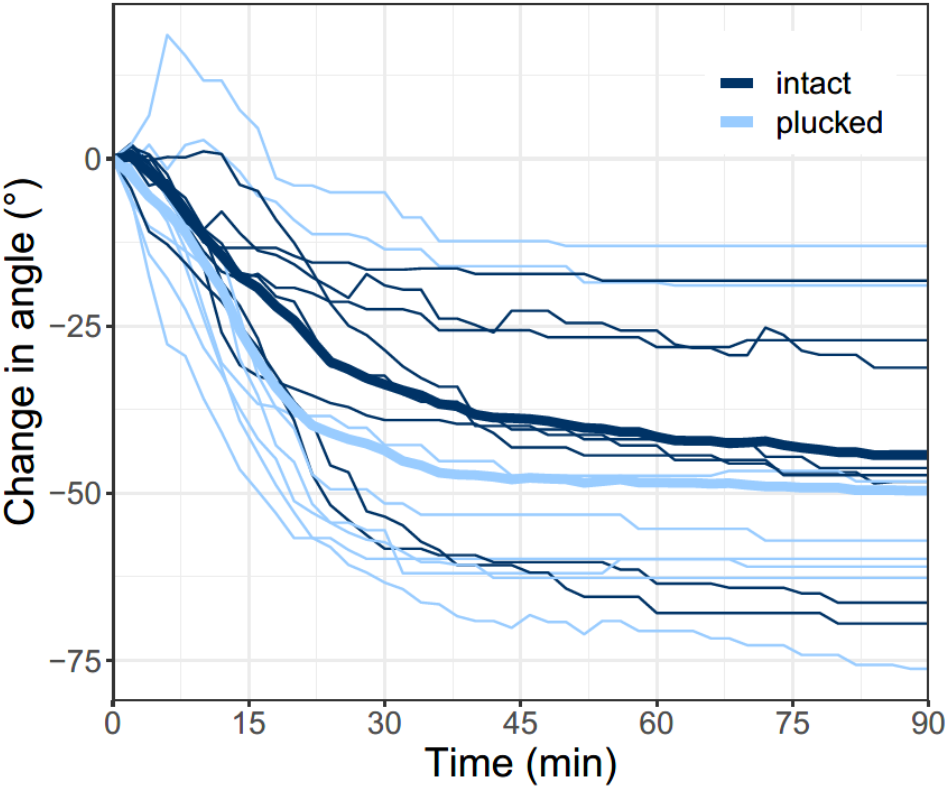
Quantification of pappus angle (the angle between outermost filaments) for plucked and intact dandelion pappi. Each sample is represented by a separate line with thicker lines indicating the mean. Dark blue indicates intact samples and light blue indicates plucked samples. n = 8 per treatment.

### The pappus actuator inhomogeneously swells to facilitate closure

As we had confirmed that the apical plate behaved as an actuator, we observed intact apical plates (actuators) and longitudinal half-sections swelling when water was added (Fig 2a-f, Supplementary Video 1). The plate is formed of cortex and epidermal cells that crease inwards towards the middle of the structure. The hairs emerge from the epidermal cells on the upper edge (Fig 2a-b). The cortical tissue is arranged around several vascular bundles and a central cavity (Fig 2c-f). Situated above all of this, is a distinct layer of tissue that originally serves as a nectary and podium for the floral organs before floral abscission and cell death occurs to form the mature pappus.

**Supplementary video 1.**
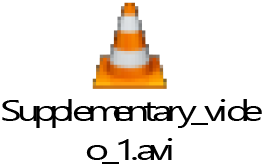
ESEM video of apical plate hydration. Chamber pressure is set to 6.3 Torr to stimulate water condensation on the sample. Once submerged with water, the chamber pressure is briefly reduced to 5.5 Torr to cause surface water to evaporate and the sample to become visible again.

**Figure 2.**
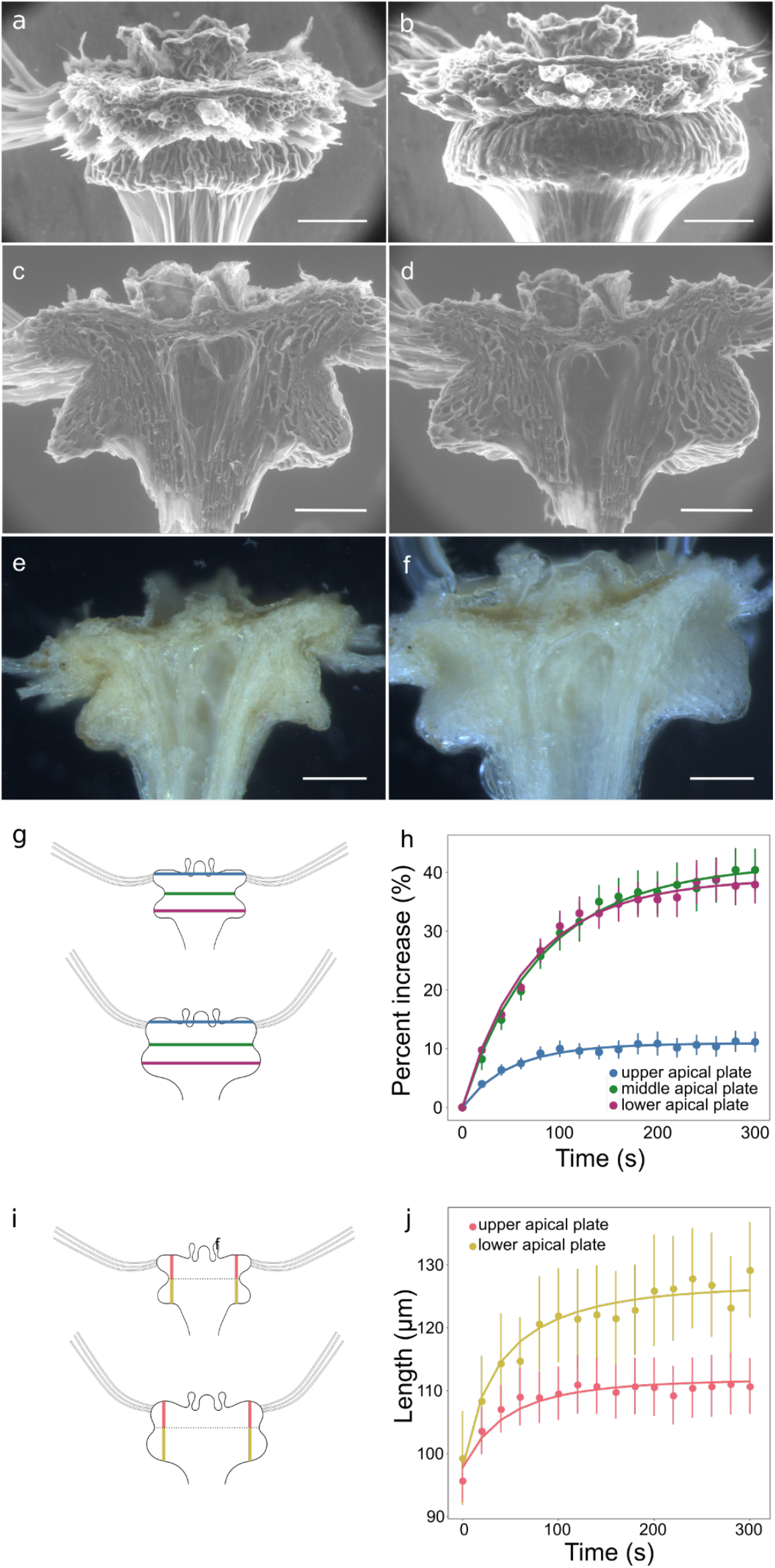
Apical plate expansion during hydration. **a-d** ESEM images of apical plate samples before, **a**, **c**, or after, **b**, **d**, hydration. **a-b** Intact sample with the majority of hairs removed. Images are representative of n=3 samples. **c-d** longitudinal half-sections. Images are representative of n = 10 samples. **e-f** Light microscopy images of longitudinal half-sections of an apical plate imaged before **e**, and after **f**, hydration. Images are representative of n = 19 samples. **g** schematic illustrating the locations of radial measurement, **h** quantification of radial expansion in half-sections imaged with light microscopy, n = 10. **i** schematic illustrating the locations of longitudinal measurement, **j** quantification of logitudinal expansion in half-sections imaged with light microscopy, n = 10. Scale bars are 100 μm.

In these circumstances we found rapid radial expansion of the actuator within 2 minutes of water addition (Fig 2e-h). Radial expansion was not uniform though as expansion was greatest between the widest points at the lower sides and the narrowest point towards the middle of the structure with an increase of around 40% in distance (Fig 2g-h). The distance between the outermost points of the floral podium only expanded by around 10% in contrast (Fig 2g-h). Longitudinal expansion was more homogeneous across the tissue (Fig 2e-f, i-j).

We observed similar results with environmental scanning electron microscopy in which the microscope chamber pressure was altered to control the condensation of water droplets on the sample (Fig 2a-d). In the dry state, most cells appear collapsed and closely packed together but the outlines of some cortical cells towards the lowermost corners of the bulging regions were visible (Fig 2c-d). Together these results indicated that the apical plate expands when wet in a heterogeneous manner.

### The actuator has distinct domains with differential cell wall composition

As the floral podium of the apical plate expanded in a different way to the lower cortical regions, we hypothesised that these tissues form a classic hygroscopic bilayer where the floral podium tissue is less able to swell than the lower cortex. This would require different material composition or properties. We found that the floral podium did not appear to be lignified in contrast to our initial expectations (Fig 3a). That layer did autofluoresce when illuminated with UV light, however, indicating phenolic compounds were likely to be present (Fig 3c). This autofluorescence increased at higher pH suggesting the presence of ferulic acid (Fig 3d,f). Using Raman spectroscopy we confirmed a high ratio of ferulic acid : lignin in this region (Fig 3g, Fig S3).

**Figure 3.**
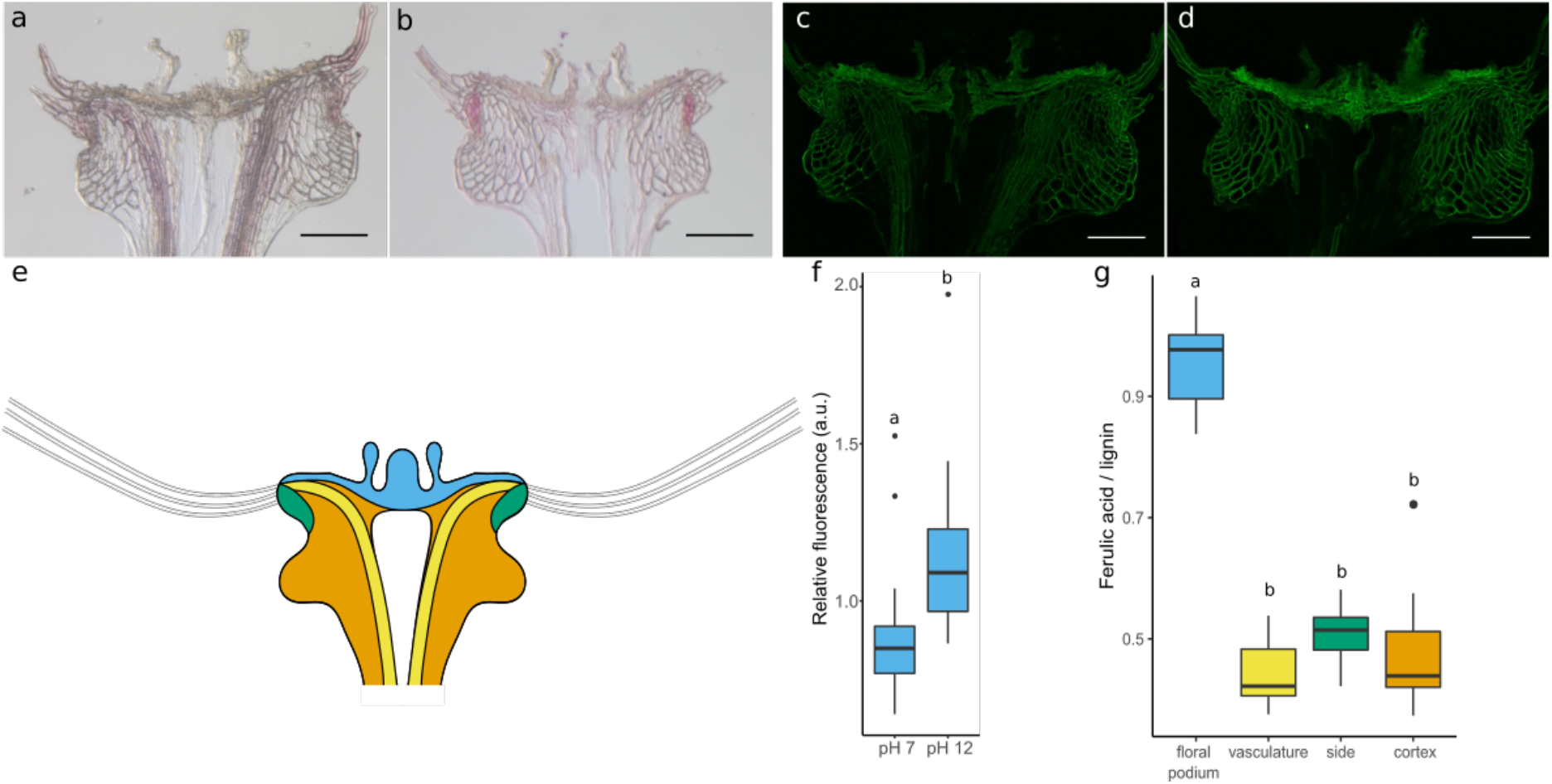
Apical plate composition. **a** phloroglucinol-HCl staining for lignin, image is representative of n = 9 samples. **b** Sudan Red 7b staining for lipids, images are representative of n = 10 samples. **c-d** UV autofluorescence at **c** ph 7, and **d** pH 12. Images are representative of n = 14 samples. **e** illustration of the different regions of the apical plate with floral podium in blue, cortex in orange, vasculature in yellow and side regions in green. **f** quantification of autofluorescence signal, n = 14 per treatment. **g** Raman spectroscopy data of ferulic acid to lignin ratio in different regions of apical plate, n = 5.

**Figure S3.**
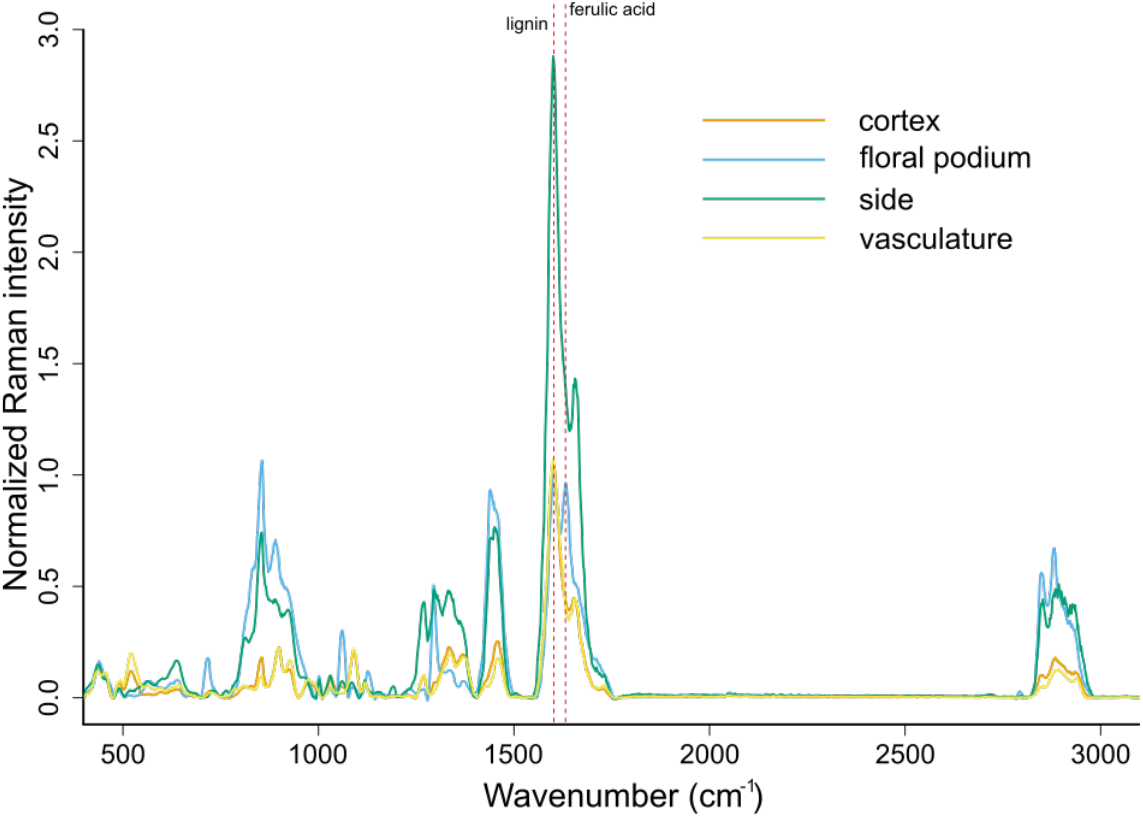
Average Raman spectra for different apical plate domains. Red dashed lines indicate the locations of peaks corresponding to lignin and ferulic acid. Data are normalized to a peak at 321 cm^−1^ arising from the CaF_2_ substrate, n = 5.

We found other regions of the apical plate with distinct cell wall compositions (Fig 3). Phloroglucinol-HCl stained most cell types but was particularly enriched in the vascular bundles indicating the presence of lignin as is common for xylem and associated fibres (Fig 3a). An intriguing lipid-rich region was also revealed around the upper sides of the cortex adjacent to the attachment site for the hairs by staining with Sudan Red 7b (Fig 3b). These data indicate that there are at least four domains of the actuator with distinct cell wall compositions or arrangements: the floral podium (blue), the vasculature (yellow), the lipid-rich sides (green) and the remaining cortex cells (orange) (Fig 3e).

### Outward radial expansion drives pappus closure

As the apical plate structure was more complex than we initially imagined, we investigated the tissue expansion in the four distinct domains (Fig 4). The autofluorescence of the phenolics was captured in longitudinal half-sections by taking high resolution z-stacks using laser confocal scanning microscopy (Fig 4a-b). The same samples were imaged when completely dry and when tissues had reached a steady state after being saturated with water. Salient landmarks, such as cell corners and small protrusions, that could be clearly identified in both dry and wet images were annotated.

**Figure 4.**
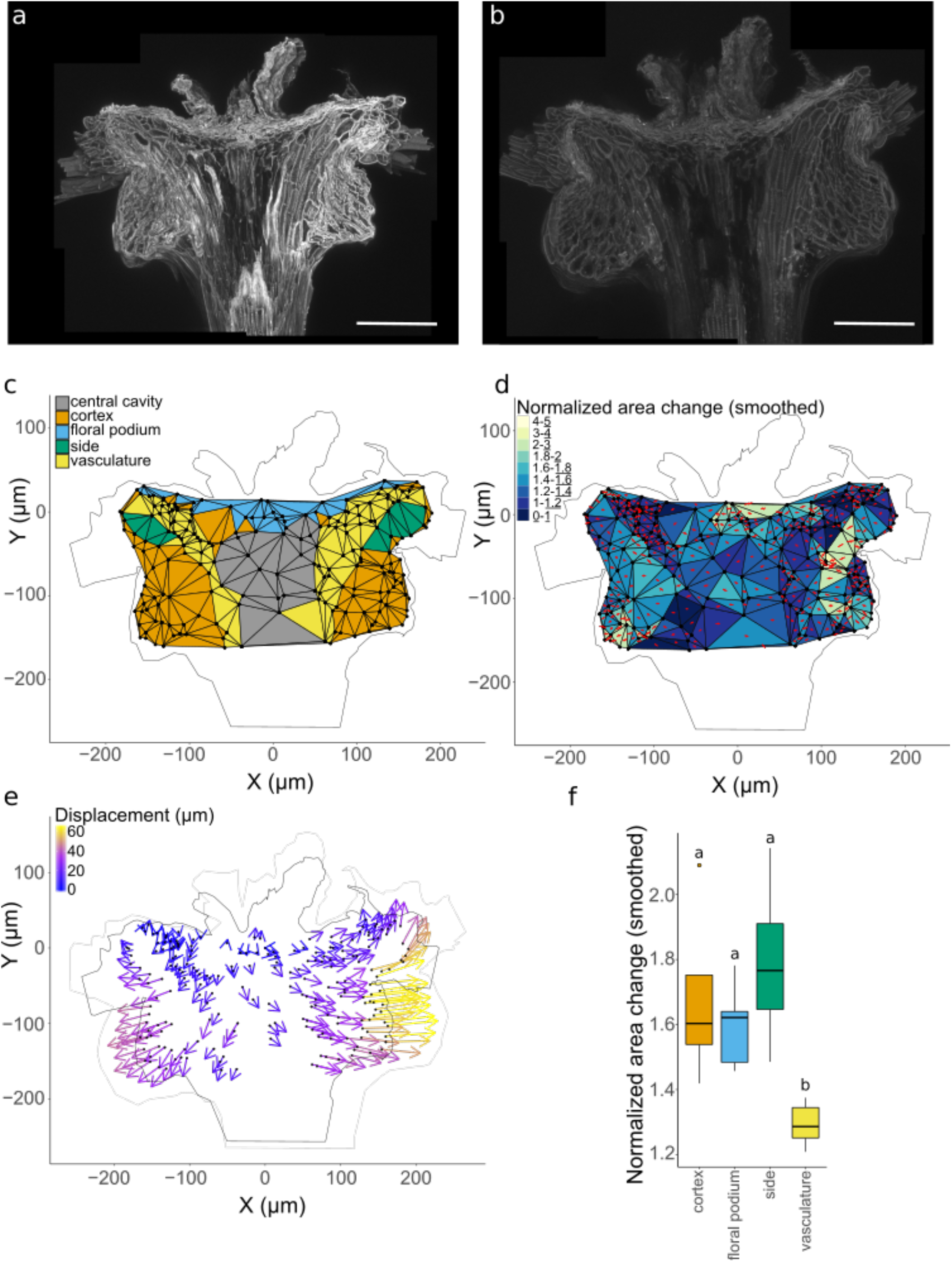
Apical plate regional expansion and displacement. **a-b** confocal images of apical plate autofluorescence in dry, **a**, and hydrated, **b** states. **c** landmarks and triangulation with regions annotated. **d** smoothed change in area of each triangle (wet area/ dry area). Red crosses indicate principal directions of expansion. **e** displacement of landmarks relative to upper central part of the apical plate. **f** change in area by region. n = 9 for all panels.

Using a central landmark close to the middle point of the floral podium as a reference, the relative displacement of landmarks was calculated (Fig 4e). The displacement map highlighted that the lower cortical regions displaced in a lateral direction from the centre and that points around the sides near where the hairs attach generally also moved outwards but also curved upwards. The vascular bundles showed limited displacement from the centre and where displacement did occur it was also largely in a radial or downward direction (Fig 4e). This tissue expansion pattern contradicts the previous hypothesis that the lower bulges of the cortex push upwards on the base of the hairs to lever them upright^20^. Instead, the radial swelling of the tissue unfolds the crease in the middle of the structure to draw the hair attachment sites outwards and rotate them into an upright position.

As point displacement reflects cumulative changes across the whole tissue, local expansion rates were also calculated to help understand the mechanism behind the anisotropic swelling (Fig 4d-e). Triangles were formed between neighbouring landmark points to allow calculation of relative expansion. These triangles were assigned an identity based on the tissue region they mostly overlapped with (Fig 4c).

Our previous measurements of radial expansion indicated minimal expansion for the floral podium tissue compared to the lower regions while longitudinal expansion was approximately uniform in the upper and lower halves (Fig 2e-j). The more detailed characterization of regional expansion demonstrated that all tissues expand to a roughly similar degree (50-90%), except for the vasculature, which shows reduced expansion (30%) (Fig 4f). This suggests that the vascular bundles may act primarily as a resisting tissue rather than the upper floral podium layer. As the vasculature is embedded within the actuator, the reduced capacity for expansion would anchor the adjoining tissues and cause anisotropic expansion overall.

### A mechanical model of pappus morphing

We hypothesized that pappus closure arises from differential water absorption or swelling properties of the different tissue regions. We also expected that the geometry and arrangement of these tissues would impact on the changing angle of the hairs. To understand how these components work together to allow pappus closure, we constructed a simplified computational mechanical model of the pappus actuator, employing Finite Element Method modelling. We considered the actuator as a two-dimensional, isotropic, linearly elastic system that undergoes shrinkage due to loss of water.

In the model, the apical plate is divided into four regions: floral podium, vasculature, sides, and cortex (Fig 5a). The tissue types were arranged according to measurements of the real dandelion pappus actuator in the hydrated configuration (Table 1). The model was allowed to shrink as if water was evaporating from the structure to generate the dry state version (Fig 5e). To account for the differences between a 2D model of a 3D phenomenon and for the simplied tissue geometry, a boundary condition was imposed that the base of the vascular bundles displaced by a fixed amount (Table 1). The angle, *θ*, between the edges of the side region and the vertical in the dry state represents the pappus holding angle and was the main prediction of the model. We compared *θ* to the equivalent measurements on the imaged longitudinal half-sections.

**Table 1.** List of measured geometrical parameters used to define the model.

| Parameter | Symbol | Value | Units | Uncertainty (%) |
| --- | --- | --- | --- | --- |
| Diameter of apical plate | D | 491.0 | $\mu\text{m}$ | 0.07 |
| Height of apical plate | H | 249.0 | $\mu\text{m}$ | 0.15 |
| Radius of curvature of floral podium | R | 363.0 | $\mu\text{m}$ | 0.14 |
| Height of floral podium | $H_{\text{pod}}$ | 38.8 | $\mu\text{m}$ | 0.25 |
| Height of side regions | $H_{\text{side}}$ | 91.4 | $\mu\text{m}$ | 0.14 |
| Width of side regions | $W_{\text{side}}$ | 47.6 | $\mu\text{m}$ | 0.14 |
| Width of vasculature | $W_{\text{vasc}}$ | 34.3 | $\mu\text{m}$ | 0.16 |
| Diameter of cavity at base | $D_{\text{cavity}}$ | 74.8 | $\mu\text{m}$ | 0.20 |
| Vascular displacement from wet to dry configuration | $d_{\text{vasc}}$ | 14.6 | $\mu\text{m}$ | 0.27 |

**Figure 5.**
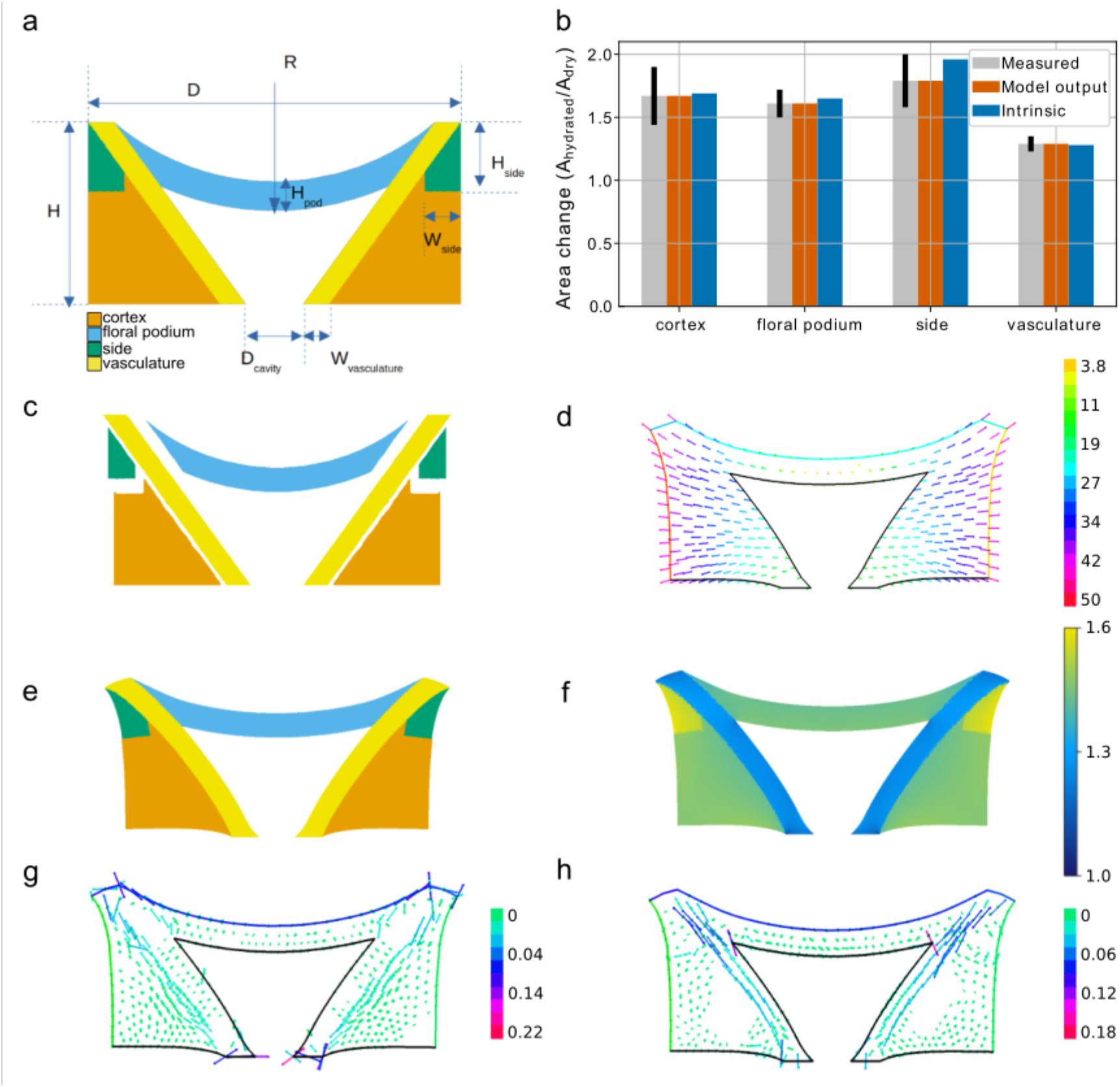
Computational modelling of apical plate behaviour. **a** geometry of the hydrated apical plate with geometrical parameters annotated. The four regions are: floral podium (blue), vasculature (yellow), sides (green), cortex (orange). **b** observed (as seen in panel **e**) and intrinsic (as seen in panel **c**) area changes by region compared to the measured area changes. **c** dry state of each region arising from different intrinsic swelling property in a hypothetical setting in which the regions are not attached to each other. **d** the displacement field relative to the center of the floral podium. **e** the dehydrated apical plate with regions annotated. Differential intrinsic swelling causes changes in shape due to the fact that they are adhered together. **f** local changes in area across the apical plate. **g** tensile stress in the dehydrated state. **h** compressive stress in the dehydrated state.

To focus on the main mechanical design features of actuation, we assigned homogenous material properties to each region taking epidermis and cortex as one tissue type (Table 2). Each region was assigned elastic properties and an intrinsic swelling property that quantifies relative changes in area upon dehydration if the region were isolated from neighbouring regions (Table 2, Fig 5c). We measured or chose plausible values for all parameters other than the four intrinsic swelling parameters (Table 1, Table 2). All geometric parameters were measured from images of apical plate sections. The cells of the apical plate appeared similar to wood cells with thickened secondary cell walls and lignification. As a result, we used a Poisson ratio typical for woody tissues (Table 2). The elastic properties were prescribed according to the measured density of cell wall material of each part (Table 2). Mechanical conflicts may arise between regions if they have different intrinsic swelling properties, leading to observed swelling that differs from intrinsic swelling (Fig 5b-c,e). The unknown intrinsic swelling parameters were fit to observations by optimising expansion in the model to the measured regional expansion from our landmark triangulation analysis (Fig 5b). We call the model together with this set of parameters the “reference model” (Fig 5).

**Table 2.** List of measured, estimated, and fitted parameters used to define material properties in the model.

| Parameter | Symbol | Origin | Value | Units | Uncertainty (%) |
| --- | --- | --- | --- | --- | --- |
| Density of cortex tissue relative to density of vasculature | $\rho_{\text{cort}}/\rho_{\text{vasc}}$ | Measured | 0.76 | none | 0.14 |
| Density of floral podium tissue relative to density of vasculature | $\rho_{\text{pod}}/\rho_{\text{vasc}}$ | Measured | 1.16 | none | 0.09 |
| Density of side tissue relative to density of vasculature | $\rho_{\text{side}}/\rho_{\text{vasc}}$ | Measured | 1.04 | none | 0.16 |
| Poisson ratio | $\nu$ | Estimated | 0.29 | none | NA |
| Swelling factor of cortex | $S_{\text{cort}}$ | Fitted | 0.46 | none | NA |
| Swelling factor of floral podium | $S_{\text{pod}}$ | Fitted | 0.44 | none | NA |
| Swelling factor of side | $S_{\text{side}}$ | Fitted | 0.57 | none | NA |
| Swelling factor of vasculature | $S_{\text{vasc}}$ | Fitted | 0.24 | none | NA |

In Figure 5b we represent the measured area changes for each region, the optimised simulated area changes as well as the corresponding intrinsic swelling. Notice the difference between the model output area changes and the intrinsic swelling per region. The regions would not change shape if they would morph independently with their own intrinsic swelling as can be seen in Figure 5c. However, due to the fact that regions are attached to one another, mechanical conflicts arise from the differential swelling between regions and gives rise to the change in shape of the structure (Fig 5d), such that the hypothetical intrinsic and actual observed area changes are different. These mechanical conflicts are visible when plotting mechanical stress patterns in the dry state (Fig 5g,h) and appear localised to vasculature and a neighbouring band along cortex and sides; vasculature is longitudinally compressed by relatively higher shrinkage in the neighbouring band, while, conversely, cortex and sides are under tensile stress parallel to the axis of vasculature due to reduced shrinkage in the vasculature.

The holding angle, *θ*, obtained as an output of the reference model is about 20° (compared to a measured *θ* = 36° ± 6.7). The displacement field relative to the centre of the podium which relates the dry to the wet state of the reference model (Fig 5e) shows similar radial displacements as was measured and shown on Figure 4e. Additionally, the local area change map that we obtain from the model (Fig 5b,f) is comparable to the measured area change map shown in Figure 4d. Therefore, the reference model sufficiently recapitulates the observed behaviour of the pappus actuator.

### The intrinsic swelling properties and dimensions of the apical plate are important for actuator function

To understand the reference model further, we carried out a one-factor-at-a-time sensitivity analysis to see which features of the model are most important for actuator behaviour (Fig 6a, Fig S4). We took the reference model and varied one parameter around its reference value while keeping all others unchanged. We monitored the relative change in the holding angle, *θ*, with respect to the relative change of the parameter in question (Fig S4). The sensitivity of *θ* to a given parameter is the ratio of these relative changes, and reflects the direct effect of this parameter on *θ* (Fig 6a). Examining the effect of each model parameter on the output *θ*, we found that the fitted intrinsic swelling capacity of each tissue had the greatest effect on holding angle (Fig 6a, Fig S4). Increasing the swelling capacity of the cortex or side regions greatly increased *θ* while the opposite was true for the vasculature and floral podium, which showed increased *θ* values when swelling capacity was decreased. For geometrical changes, the sensitivity analysis highlighted the overall dimensions in the horizontal and vertical directions (*D* and *H*) and the radius of curvature of the floral podium (R) as having substantial effects on the holding angle change (Fig 6a, Fig S4).

**Figure 6.**
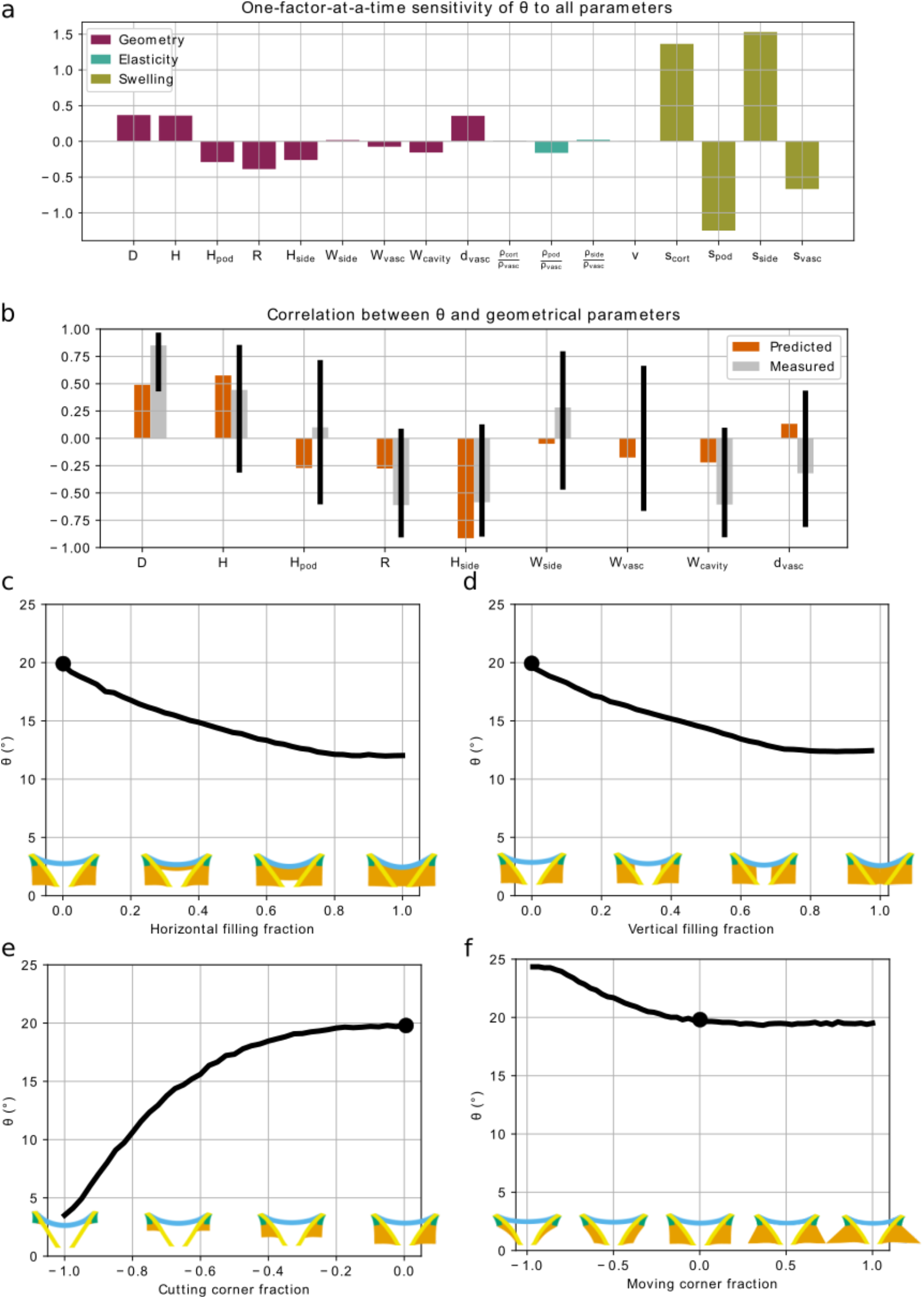
Predictions of the output holding angle (*θ*) resulting from different perturbations of the model. **a-b** perturbations of the parameters. **a** the partial sensitivity of *θ* to input parameters obtained by an OAT (one at a time) sensitivity analysis. **b** the computed (predicted) correlation of geometrical input parameters to *θ*, when correlation between input parameters is taken into account, are compared to the measured correlation coefficients. Black bars denote 95% confidence intervals around the mean measured value. **c-f** perturbations of the geometry of the reference model (marked by a black circle) by adding (positive values) or removing (negative values) cortex material. **c** the central cavity is filled horizontally. **d** the central cavity is filled vertically. **e** cortex material is removed from the corner of the actuator vertically. **f** cortex material is removed from or added to the actuator by moving the corners horizontally.

**Figure S4.**
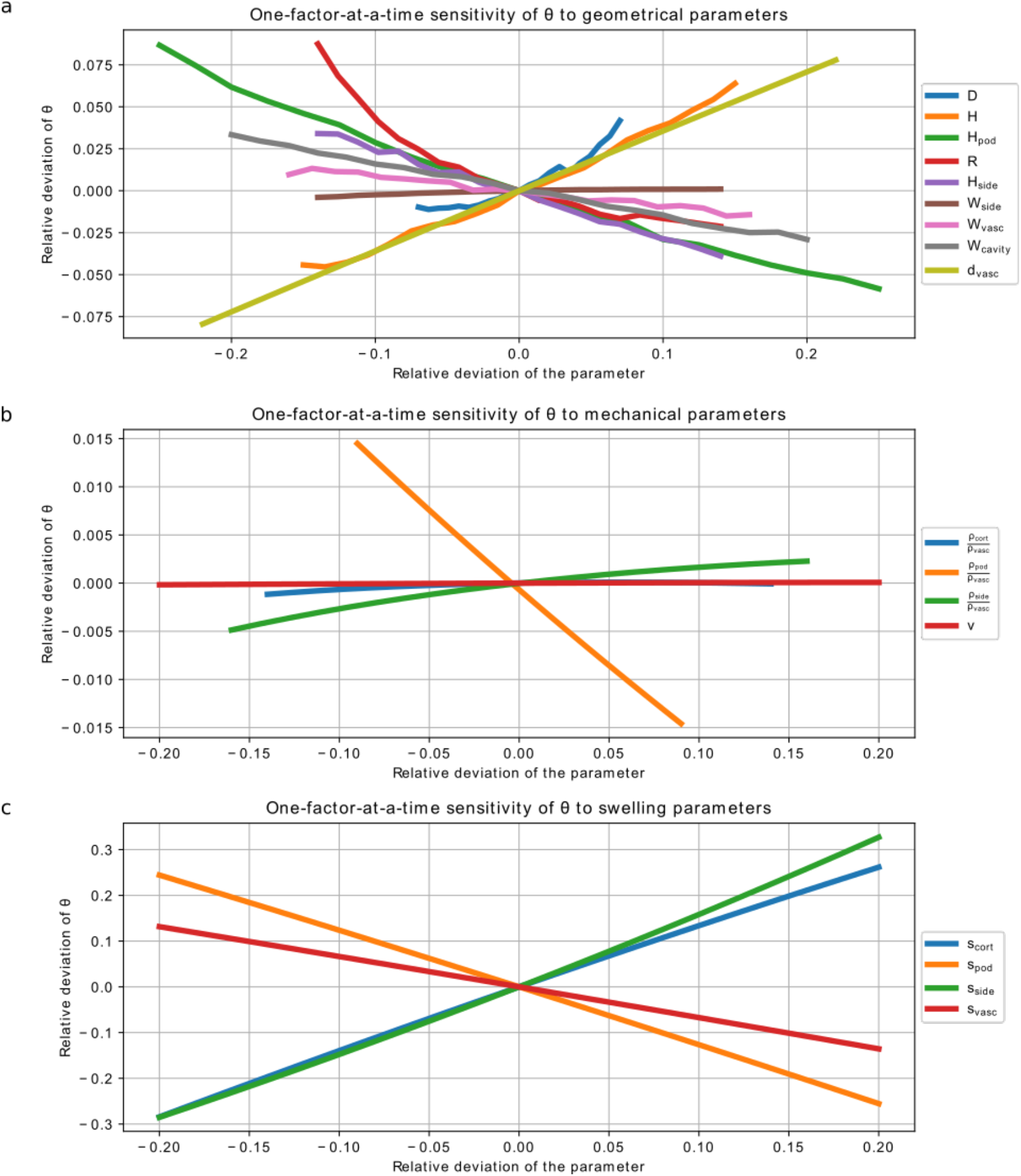
One-factor-at-a-time sensitivity analysis across the range of parameter variations. **a** Sensitivity analysis of geometrical parameters. **b** Sensitivity analysis of mechanical parameters. **c** Sensitivity analysis of swelling parameters.

In theory, our model predicts how angle change varies with each of the actuator geometrical parameters. However, these geometrical parameters are not independent and co-vary in biological specimens (Fig S5). To test the validity of our model, we used the sensitivity analysis to compute the correlation between *θ* and geometrical parameters given the measured natural correlations between geometrical features observed in our specimens (Fig 6b, Fig S5). The predicted correlation between *θ* and geometrical parameters was compared with measured correlations between these features. The confidence intervals were generally large for most measured correlations, but for those that were statistically significant, the predicted correlation coefficients fell within the expected range for all but one (*H_side_*) parameter (Fig 6b). We surmised that the cause of this large negative correlation between *H_side_* and *θ* is due to the way in which *θ* is measured. Note that *θ* was determined as the angle between the vertical, and the line formed between the upper corner of the apical plate to lateral edge at the lowest point of the side region, which has height *H_side_*in the dehydrated state. Therefore, when the parameter *H_side_* was varied in the sensitivity analysis, the line giving the definition of the holding angle *θ* also changed. We therefore believe that the large negative correlation between *H_side_* and *θ* in the sensitivity analysis is largely due to the changing location of this measurement line.

**Figure S5.**
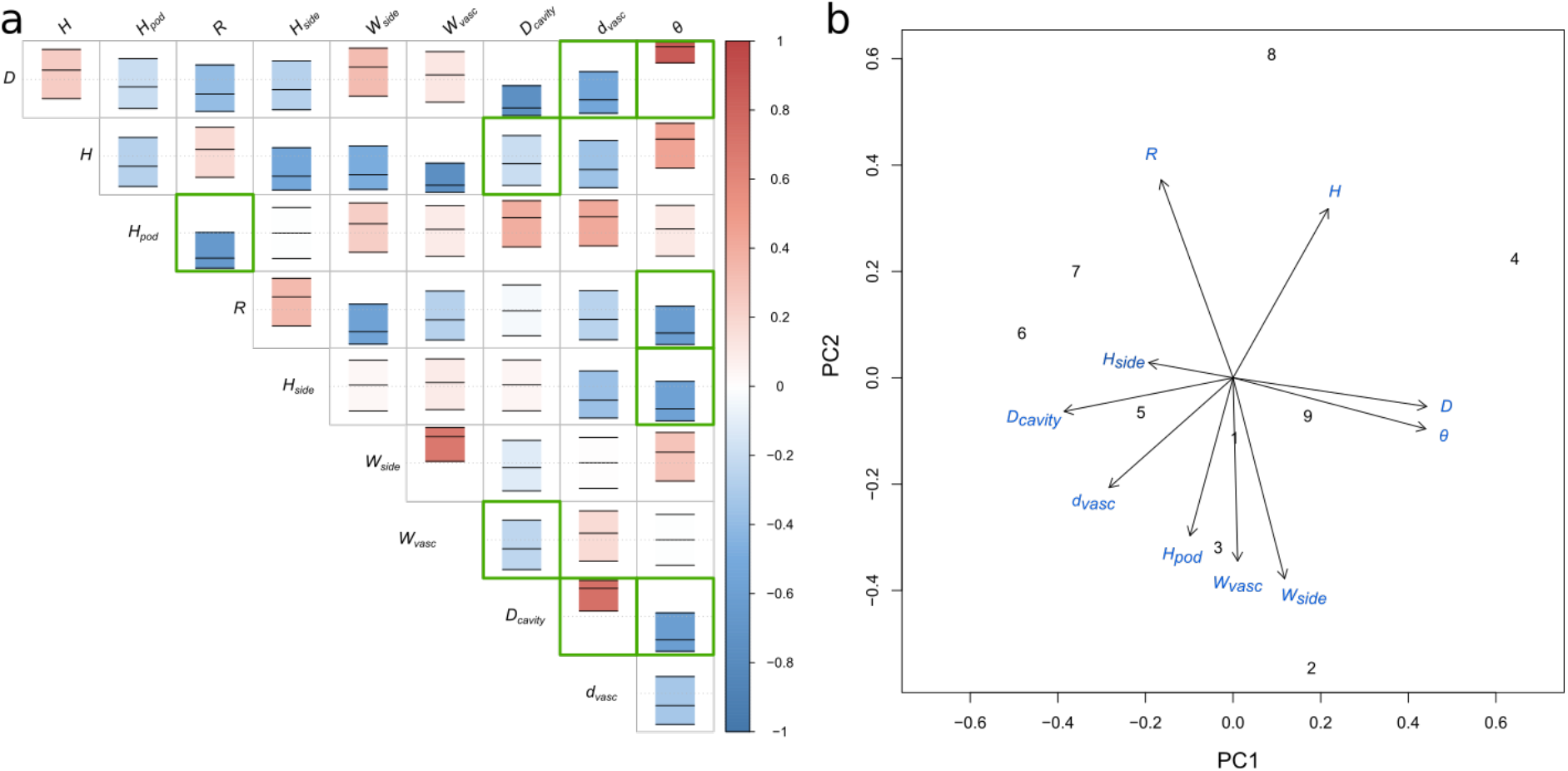
Relationship between observed geometrical features. **a** correlation matrix illustrating the relationship between observed geometrical parameters. The colour scale and central bar of each box indicates the correlation coefficient, outer bars give the 95% confidence interval limits. Green boxes indicate a statistically significant result at p < 0.1. **b** biplot of the first two principal components from a principal component analysis of geometrical parameters. n = 9.

As in the one-factor-at-a-time sensitivity analysis, the sensitivity analysis accounting for natural co-variations highlighted the importance of *H* and *D* in positively regulating *θ* (Fig 6b). *H_pod_* was less significant in this context, while *H_side_* appeared much more important for determining *θ*. In conclusion, this analysis predicted correlations between *θ* and geometrical parameters, in fair agreement with experimental measurements (Fig 6b).

### The material properties of each domain affect actuation

To understand further the extent to which material properties affected actuator behaviour, we created hypothetical scenarios in which certain regions were made of alternative tissue types (Fig S6). We found that substituting the material of each region with cortex-like material generally reduced the magnitude of the holding angle, *θ* (Fig S6a). Despite this, we observed that the model was still quite robust and achieved moderate angle changes (>14°) when all regions contained cortex-like tissue, provided the vasculature retained its original properties of higher stiffness and minimal swelling capacity (Fig S6a). When the side regions were the only contrasting tissue to cortex, only a small *θ* was observed but when combined with vasculature in its natural state, an enhanced *θ* was observed comparable to the reference model (Fig S6a). This indicates that the distinctive vasculature material properties (low swelling capacity and moderately high stiffness) relative to the other regions are most important for actuator behaviour. In this case, substituting the floral podium for cortex tissue had no substantial effect on the resulting holding angle (Fig S6a) probably because the material properties (in particular the swelling factor) of these two regions are quite similar (Table 2).

**Figure S6.**
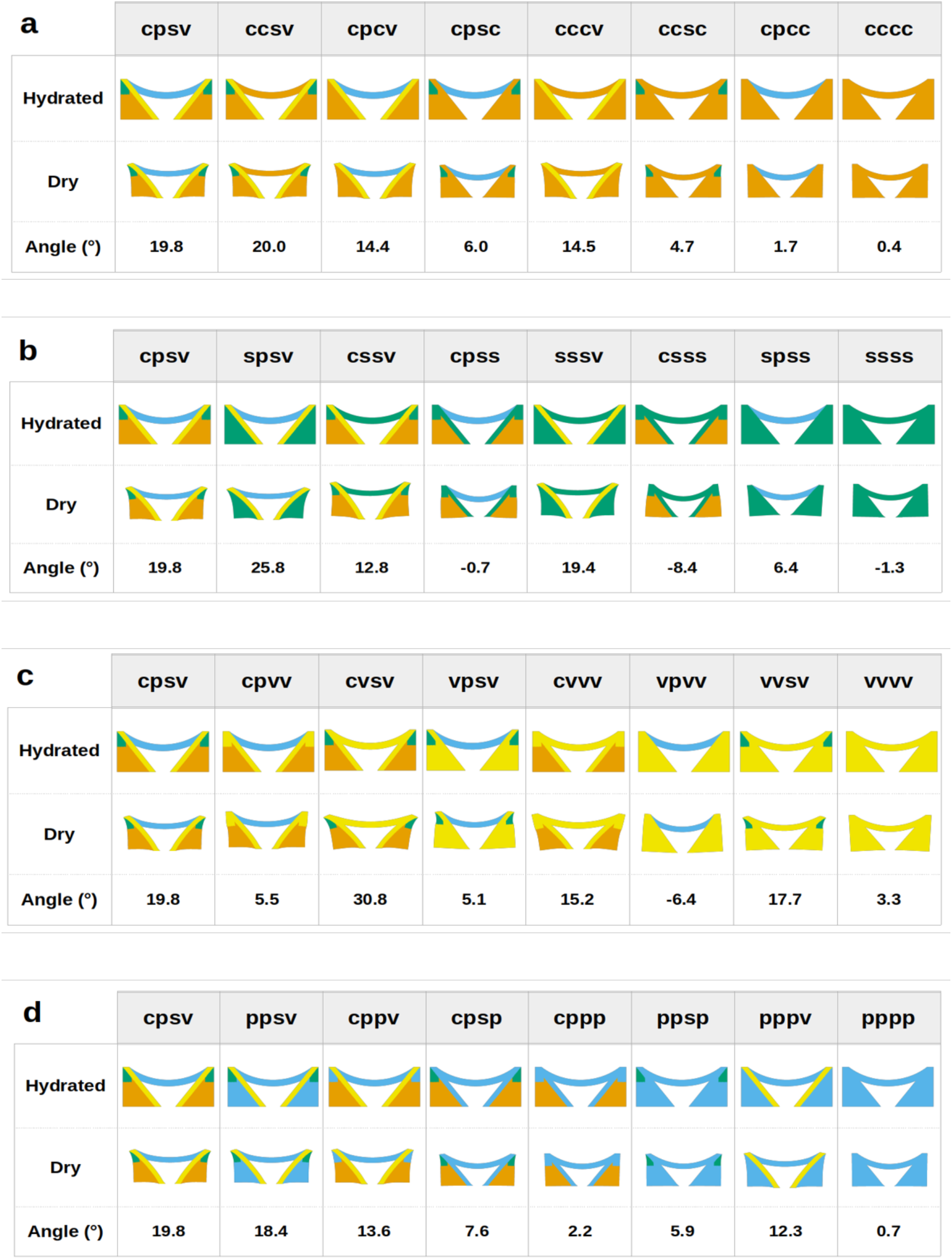
In silico mutations. Each region is substituted with alternative tissue types. Tissue types are denoted by letters (c=cortex, p=floral podium, s=side, v=vasculature), and the different geometrical regions are marked by the order of letters in the word designing the mutant (1^st^ letter=cortex region, 2^nd^ letter=floral podium region, 3^rd^ letter=side region, 4^th^ letter=vasculature region in the reference model). This way the reference model is denoted by the word “cpsv”, and a mutant in which the vasculature region is turned into cortex is denoted “cpsc”. a: substitutions with cortex-like tissue. b: substitutions with side-like tissue. c: substitutions with vasculature-like tissue. d: substitutions with floral podium-like tissue.

We then substituted each region with material with side region-like properties (Fig S6b). Making the vasculature even more swellable than the cortex (swelling factor 0.57 vs 0.46, Table 2) entirely abolished the actuator function and if, in addition, the floral podium was also substituted with side material, the actuator began to invert and generate negative *θ* values (−8.4°) (Fig S6b). Substituting only the floral podium for the side material that swells more also reduced the actuator function, though to a lesser extent (12.8°). This indicates that though primarily the vasculature properties must contrast with the cortex, the floral podium also plays a resisting role. Interestingly, the actuator function could be enhanced by replacing cortex with side material indicating that the actuator may not be perfectly optimised for maximising holding angle changes (Fig S6b).

We also attempted to substitute regions with the vasculature tissue type to assess the effect of reducing swelling capacity and, for some regions, increasing stiffness (Fig S6c, Table 2). Applying this tissue type to either cortex or side regions caused only a modest reduction in *θ* indicating that a structure possessing either contrasting tissue type adjacent to the vasculature can be almost sufficient (Fig S6c). However, this relied on also having a floral podium that could not swell easily (i.e. replaced with vasculature material). Without this restraint, the actuator almost completely failed. If the floral podium was replaced with vasculature but the other regions remained normal, the largest holding angle changes were observed compared to any other scenario including the reference model (Fig S6c).

Together, these data suggest that for the apical plate to function best as a hygroscopic actuator, the vasculature and floral podium must be unable to swell easily while the cortex must easily swell. Highly swelling side regions can enhance the behaviour of the actuator (though increasing the size of that domain can be detrimental) but are not essential to broadly capture the actuating function (Fig S6).

### The arrangement of tissues around the central cavity is essential for changing holding angle

To explore the importance of the observed apical plate geometry, we made some modifications to the geometry of the computational model and assessed the predicted holding angle, *θ* (Fig 6c-f). We found that the central cavity between the vascular bundles and beneath the floral podium was necessary to allow substantial angle changes (Fig 6c-d). Progressive filling of the cavity with cortical tissue either laterally or longitudinally correspondingly reduced *θ* (Fig 6c-d)

We also test the role of the cortex tissue surrounding the vasculature (Fig 6e-f). We found that altering the size of the lower bulges in the lateral direction had very little effect on *θ* (up to 5° difference from the reference model) and reducing the tissue to only a thin layer around the vasculature was actually able to slightly increase *θ* (Fig 6f). In contrast removing the cortex tissue longitudinally caused a dramatic reduction in the angle change (a decrease of 14° from the reference model) (Fig 6e). This indicates that the quantity of cortex tissue present around the vasculature is not important, but that the connection between the side regions and the whole length of the vasculature must be maintained (Fig 6e). Together, these results indicate that the geometry we observe is almost optimal for pappus closure in comparison to the majority of modifications we made. They also highlight a conical vasculature-cortex bilayer structure at the core of the actuator’s functioning: both removing the outer cortex layer adjacent to the vasculature (Fig 6e) and adding a third internal cortex layer in the central cavity (Fig 6c-d) considerably impairs the morphing capability of the structure.

## Discussion

Our data indicate that the balance of resisting and swelling tissues is carefully arranged in the dandelion actuator to allow functional angle changes (Fig 3, Fig 6, Fig S6). The vasculature and floral podium together provide a resisting framework anchoring the surrounding swellable tissues that provide the majority of the swelling motion. Additionally, the central cavity is essential to provide space for the other regions to contract into when drying (Fig 6c-d).

This arrangement allows precise radial swelling that is not seen in other hygroscopic movements. While there are other systems that involve the bending of filamentous structures that are radially arranged (such as in the unfurling of *S. lepidophylla* stems^17^ and in the bending pedicels of carrot umbels^15^), these rely on separate bending of each stem individually. For the smaller dandelion pappus with exceptionally fine hairs^26^, it is possible that a single actuating structure is more efficient and allows better co-ordination of motion. This might prevent tangling or breakage of hairs if they were misaligned during bending. We expect that this is not unique to the dandelion as many other Asteraceae exhibit hygroscopic opening and closing of their pappi^19,21^. Additionally, many plant structures are inherently radial so it is likely that a pre-existing structural arrangement of vascular fibres and other tissues could be easily selected for during evolution to facilitate biomechanical movements.

The hygroscopic plant movements that have so far been described involve a combination of structural and compositional features of cell walls that result in either bending, twisting or coiling motions. These typically rely on a bilayer structure consisting of a swelling tissue or cell wall compartment and an adjacent resisting element. The dandelion apical plate can be considered as a variation of bilayer structure since swelling regions must be connected to regions that do not swell. The radial motion, however, means that a more complex structure is required consisting of at least three tissue types (Fig 3). In our model, structures with only two tissue types performed less well than the reference model (Fig S6). Even in cases where two tissue types gave a *θ* value comparable to the reference model (Fig S6b – sssv, S6c – vvsv), there were circumstances where adding a third tissue type could further enhance *θ* beyond the level of the reference model (Fig S6b – spsv, S6c – cvsv).

Assigning reduced swelling properties to the floral podium enhanced the pappus holding angle changes in the model (Fig 6, Fig S6) (though note that increasing the height or density of the podium reduces *θ* (Fig 6a)). This indicates that having a three-part structure in which a bilayer is combined with another resisting material on an intersecting plane further constrains some of the swelling to enhance anisotropic motion. This is more reminiscent of the cup-shaped thickening in the individual cells of fern sporangia that collectively resist bending during dehiscence and allow directed opening of the spore capsule^27^. It is possible that in the 2D model, the floral podium acts primarily to connect the tissues and keep the two sides separate. A 3D computational model may change this requirement as the remaining tissue would be radially connected.

Our data and modelling give some hint as to the mechanism behind the differential swelling properties of each region. The strong lignification of the vascular tissue (Fig 3a) suggests that these cells are highly hydrophobic, which corresponds to the minimal expansion observed. Ferulates in the floral podium (Fig 3c-d,f) also indicate a hydrophobic material though it is also likely that high density of cell walls (Table 2) causes that layer to act as a partially resistive tissue. Though the cortical cells do appear to be somewhat lignified (Fig 3a,c), the cells are larger with significant spaces between them (Fig 2c-f, Table 2). This may mean that for a given proportional swelling of those cell walls there are greater absolute levels of expansion at the tissue level compared to smaller, denser cells in other tissues.

The side regions contain a lipid-rich material (Fig 3b), which might be suberin. While suberin is classically associated with the hydrophobic Casparian strip of the root endodermis and abscission zones^28^, the side cells of the dandelion apical plate expand considerably with wetting. While this appears contradictory, the relative hydrophobicities of cell wall materials are not well understood and in fact Casparian strip water impermeability appears to rely more on ferulate components than on waxes^29^. Another possibility is that the lipid component maintains cells in a flexible but hydrophobic state such that they are able to expand and hold water within them. This would suggest that they act more like a balloon holding fluid inside than a sponge that absorbs water into its material structure. This may explain why high relative humidity has only a limited effect on pappus closure and that liquid water is generally required to achieve complete morphing^22^. An analogous phenomenon has been demonstrated for the ice plant seed capsules that require liquid water for opening behaviour^30^.

A feature of other hygroscopic systems is that they frequently rely on inherent anisotropic swelling of the cell walls by orientating cellulose microfibrils^8,9^. In the dandelion apical plate, we cannot rule out intrinsic swelling anisotropy as a possibility. However, we find from our computational model that intrinsic anisotropy is not necessary for a functional model. Simply juxtaposing tissue types with differential isotropic swelling capacities and stiffnesses that arise solely from cell density recapitulates the behaviour of the actuator (Fig 5). While there are some small differences between the model and experiments, this might be due to 2D modelling of a 3D phenomenon and even the 20° change in angle observed in the model would substantially impact flight behaviour^22^.

The hygroscopic actuator underlying dandelion pappus morphing is made of non-living cells, whose cell walls have differential water-dependent expansion and drive the movement of the pappus hairs. It is a previously uncharacterised type of biological hinge, which is a radial, tubular actuator that can coordinate collective movement of hairs positioned on a planar disk like an umbrella at the periphery. The dandelion actuator has design elements resembling bilayer structures, yet it is distinct from planar or cylindrical bilayer actuators that bend or fold, which have been well explored in biomimetic soft robotics. From intestines and bladder to sea anemones, tubular actuators are abundant in living systems and represent a next frontier for biomimetic soft robotics. The simple design requirement of the dandelion actuator makes it likely possible to be fabricated at micro-scale or smaller, and may be applied to the creation of novel animated functional materials.

## Acknowledgements

We thank Jim Buckman at Heriot-Watt University for use of the environmental scanning electron microscopy facility and Erika Kroll for assistance with pappus imaging. We also thank staff of the Nikolai Lobachevsky Library of Kazan Federal University for access to a scanned copy of the paper by Taliev in ref ^19^. This project was supported by a Leverhulme Trust grant (RPG-2015-255). NN is supported by a Royal Society University Research Fellowship (UF140640 and URF\R\201035) and MS is supported by Leverhulme Trust Early Career Fellowship (ECF-2019-424).

## Author Contributions

MS, AK, EM, AB, and NN designed the experiments, MS, AK, and SB conducted the experiments and analyses, all authors contributed to interpretation of the results, MS, AK, AB, and NN drafted the paper, and all authors revised the manuscript.

## Materials and Methods

### Plant material

*Taraxacum officinale* samples were collected and grown as described previously^26^. All samples originated from the same original individual and were grown in the greenhouse for two generations. The offspring of these were used for experiments. Diaspores used were assumed to be genetically identical as this subspecies reproduces apomictically.

### Moisture chamber imaging

A bespoke moisture chamber was set up to release small mist droplets into a air-tight box containing dandelion fruits. This was assembled as described in ref ^22^ with an ultrasonic humidifier for water release and small USB microscopes (Maozua, USB001) for imaging. A datalogger (Lascar Electronics, EL-GFX-2) was included in all experiments to monitor temperature and relative humidity. For all experiments, moisture was added to the chamber for 20 minutes and pappi allowed to absorb water and equilibrate with their surroundings within the sealed chamber. Images were captured at the start of the experiment before water was added and after three hours.

For the apical plate blocking experiments, methacrylate nail polish was carefully applied using a sewing pin and subsequently cured by exposing to UV light. For blocking of the lower side of the apical plate the hairs of dandelion pappi were temporarily tied loosely together using cotton thread to allow access. Control pappi had hairs temporarily tied together but no nail polish applied.

To test the effect of increasing hair spacing, pappi were either left intact as a control treatment or had the majority of hairs carefully removed using fine forceps. Two hairs were allowed to remain that were approximately opposite one another.

### Microscopy and histology

For longitudinal half-sections, dandelion pappi with hairs mostly removed were directly embedded into paraffin wax without any fixing or infiltration. A microtome was used to slice away material and the cut surface examined periodically under a microscope. Once the central cavity and the bulge at the centre of the floral podium (the stylopodium) were visible, the apical plate was considered to be cut in half. The majority of the wax was cut away manually and the remaining half section was briefly submerged in Histoclear to dissolve any remaining wax.

To visualise cell wall materials using cell wall stains and autofluorescence, dandelion pappi were progressively infiltrated with water followed by ethanol, Histoclear and paraffin wax. Longitudinal sections of 10 μm thickness were made using a microtome and sections were dewaxed using the inverse infiltration procedure. For lignin staining, 2 parts of 3% (w/v) phloroglucinol dissolved in absolute ethanol and combined with 1 part concentrated HCl according to ref ^31^. Sections were imaged immediately using a stereomicroscope. Lipid staining was carried using 0.1% (w/v) Sudan Red 7B dissolved in PEG-300 and combined 1:1 with 90% glycerol according to ref ^32^. Staining was carried out for 16 hours before washing with water and mounting in glycerol and imaging using a stereomicroscope. Autofluorescence images were acquired using a confocal microscope with an excitation wavelength of 405 nm and emission acquisition from 480 – 600 nm. For ferulic acid identification, paired sections were used in which two adjacent longitudinal slices close to the medial section of each apical plate sample. These were imaged using a 10x objective at either pH 7 (distilled water) or pH12 (0.01M KOH) for 15 mins^33^.

To image apical plate expansion, apical plate half sections were lightly glued to a glass slide with a small dot of epoxy resin and a coverslip overlaid. Images were taken of dry samples and when wet by pipetting distilled water underneath. For time course imaging, a stereomicroscope was used and images captured every 20 seconds. For landmark annotation, a 63x objective was used to image autofluorescence of the cut face of the samples. Tiled z-stacks were acquired and maximum projections were later stitched together of each sample when dry and wet after allowing the tissue to reach a fully expanded hydrated state (around 30 minutes).

Apical plate expansion was also imaged in an environmental scanning electron microscope (ESEM) (FEI Philips XL30). Half-section samples were prepared as described above and placed onto an adhesive carbon pad to mount onto the ESEM stage. The Peltier stage was maintained at 5°C and dry samples imaged by maintaining chamber pressure at 5 to 5.5 Torr. To encourage water condensation, the chamber pressure was increased to 6.7 Torr. Once the sample became fully submerged in water the surface was no longer visible so after allowing sample expansion to occur, the pressure was decreased again to 5.5 Torr. This caused water to evaporate and images of hydrated tissue were rapidly acquired.

### Raman spectroscopy

Longitudinal 10 μm sections (see Microscopy and histology) were mounted on CaF_2_ cover slips, submerged in distilled water, and imaged with a Renishaw InVia Raman microspectrometer using a 63x dry objective. A 785 nm laser beam was used for excitation at a laser power of 100% and exposure time of 10 s. Spectra were obtained from at least 20 points per region (floral podium, vasculature, cortex and sides) for each sample and 5 samples from different individual diaspores were used.

A rolling ball baseline correction was applied to all spectra with a baseline identification window of 60 and a smoothing window of 3. Intensity values were normalised to the CaF_2_ substrate-derived peak at 321 cm^−1^. To calculate the ratio of ferulic acid to lignin, the intensity at 1631 cm^−1^ corresponding to ferulic acid was divided by the intensity at 1602 cm^−1^ corresponding to lignin ^34,35^. Tukey’s honest significant difference test was used to detect statistically significant differences between regions.

### Image analysis

To measure pappus angle from the moisture chamber imaging experiments, images identity was blinded and images were randomised before measurement. Using Fiji, the angle between the outermost filaments (excluding those that were not in focus) was measured. Measurements of cell and tissue sizes and lengths were carried out using line or polygon measurement tools in Fiji.

Autofluorescence was quantified by measuring the mean gray value in small regions of the floral podium from confocal images. For each sample, fluorescence was normalized to the autofluoresence derived from the pappus hairs at pH 7.

For landmark displacement and relative expansion measurements, z-stacks of confocal autofluorescence images were used as described above. Maximum projections of each z-stack were obtained and images were stitched together using the Fiji plugin, Pairwise Stiching^36^. The polygon measurement tool was used to generate an outline of both dry and wet apical plates. Matched landmarks for dry and wet images of each sample were manually annotated. These consisted of salient features, such as cell corners, ridges and holes. For each sample, one landmark was selected close to the centre around the junction between the floral podium and central cavity to act as a reference point. Displacement of each landmark was calculated relative to this central point.

For local expansion rates, a Delaunay triangulation was mapped onto the landmarks of the dry state samples using R package deldir^37^. Triangles that were not fully enclosed by the overall outline of the dry sample were excluded. The area of each triangle was calculated and normalized area change calculated by dividing the area of a triangle when wet by its area when dry. The data were smoothed by calculating the arithmetic mean of the normalized area change for each triangle and its adjoining neighbouring triangles. Principal orientations of strain was calculated for each triangle and annotated as a cross scaled to 3 times larger than the original values for improved visibility^38^.

Regions of the apical plate were designated by manually outlining the floral podium, vasculature, side regions and central cavity on the original images using the Fiji polygon tool. Triangles in the dry state that overlapped at least 40% with any of the dry state regions were assigned to those regions and all others were designated as cortex triangles. Mean area changes were calculated for each region for each sample and these values used to generate boxplots and for statistical analysis.

For a very small number of triangles the nonuniform expansion process rearranged nearby landmarks relative to one another such that triangles then appeared to overlap in the wet state. As it is not physically possible for cells to actually overlap in a connected cellular structure, these triangles were omitted from the analysis and comparisons between regions.

Geometrical measurements of the apical plate were obtained from images and used as inputs for the computational model. Measurements of the sizes and geometry of regions were taken from the wet-state confocal images of apical plate half-sections using Fiji. For some parameters such as the vascular displacement (d_vasc_) and holding angle (θ), measurements were taken from consistent points on the dry and wet images and differences calculated. For region density measurements, 10 μm sections stained with ruthenium red gave consistent red staining across all cell types. These images were thresholded using the automatic colour thresholding in Fiji, converted to a binary image and the ratio of stained to unstained pixels used to calculate the density of cell wall material in manually selected rectangular areas of each region.

### Statistical analysis

Student’s t-test was used to detect differences in cell area change in the wet and dry states. Tukey’s honest significant difference test was used to detect statistically signficant differences in pappus angle and for comparisons of quantitative data between regions of the apical plate.

Nonlinear mixed effects models were fitted to the time course of radial expansion using R package ‘lme4’^39^. A negative exponential function was fitted using a modification of the ‘SSAsymp’ function in lme4. Time and measurement location were fixed effects while sample ID was a random effect. A model was selected based on comparing AIC. The optimal model had slopes (natural logarithm of the rate constant) and asymptotes varying according to sample ID and measurement location with the intercept constrained to 0. This model performed better than versions in which the slope, asymptote or both only varied by sample and not measurement location.

The correlation coefficient was calculated for geometrical parameters using the R package ‘corrplot’^40^ and the biplot of principal components plotted from a principal component analysis computed in R.

### Modelling

A detailed description of the model is included in the supplementary materials and a brief description provided here.

The hydrated state of the actuator is corresponds to the geometry of the living tissue with turgor pressure removed. We consider that in this case, all stresses are relaxed. Therefore, an important assumption of our model is that the hydrated state of the actuator is completely stress-free. Starting with this stress-free state, we modelled the actuator as a two-dimensional, isotropic, linearly elastic system that undergoes shrinkage due to loss of water. The corresponding partial differential equation corresponds to stress balance, with a source term associated with shrinkage. These equations are solved using FreeFem++^41^.

The starting hydrated geometry of the system is divided into 4 regions, with an arrangement and dimensions that follow experimental measurements. The displacement of vasculature at the basis of the actuator was prescribed according to experimental measurements. All regions have the same Poisson ratio. The podium, vasculature, and sides are characterised by the ratio of their modulus to that of cortex, which was inferred from density of cell wall material in experimental data. Each region has an intrinsic swelling parameter, which cannot be directly measured. Swelling parameters were determined by minimizing the difference in observed shrinking between model and experiments. We call the model, together with the measured and fitted parameters, the reference model.

The main prediction of the model is the holding angle, *θ*. A higher value of *θ* indicates a more substantial pappus opening during dehydration indicating more efficient functioning of the actuator. Model sensitivity to a parameter was assessed from the derivatives of the angle *θ* with respect to this parameter around neighbouring values to the reference value. Correlations between *θ* and geometric parameters were predicted by combining the sensitivity values with the correlation matrix of geometric parameters (see supplementary materials for details).

## Data availability

Data will be made available by the corresponding authors on reasonable request.

